# Coordinated prefrontal state transition leads extinction of reward-seeking behaviors

**DOI:** 10.1101/2020.02.26.964510

**Authors:** Eleonora Russo, Tianyang Ma, Rainer Spanagel, Daniel Durstewitz, Hazem Toutounji, Georg Köhr

**Author notes:** These authors contributed equally (Hazem Toutounji, Georg Köhr).

## Abstract

Extinction learning suppresses conditioned reward responses and is thus fundamental to adapt to changing environmental demands and to control excessive reward seeking. The medial prefrontal cortex (mPFC) monitors and controls conditioned reward responses. Using *in vivo* multiple single-unit recordings of mPFC we studied the relationship between single-unit and population dynamics during different phases of an operant conditioning task. To examine the fine temporal relation between neural activity and behavior, we developed a model-based statistical analysis that captured behavioral idiosyncrasies. We found that single-unit responses to conditioned stimuli changed throughout the course of a session even under stable experimental conditions and consistent behavior. However, when behavioral responses to task contingencies had to be updated during the extinction phase, unit-specific modulations became coordinated across the whole population, pushing the network into a new stable attractor state. These results show that extinction learning is not associated with suppressed mPFC responses to conditioned stimuli, but is driven by single-unit coordination into population-wide transitions of the animal’s internal state.

## INTRODUCTION

The medial prefrontal cortex (mPFC) plays a key role in numerous behaviors and cognitive functions, including action control, attention, cognitive flexibility, decision-making, and reward learning (Euston et al., 2012; Ridderinkhof et al., 2004; Stoll et al., 2016). Reward-driven learning guides our daily life activities and requires associating specific cues and environments with reward. The resulting conditioned reward-seeking responses are monitored and controlled by the mPFC (Botvinick et al., 2019). Lack of such cognitive control can lead to maladaptive behaviors, such as substance abuse or perseveration in reward-seeking responses when reward availability has ceased. Extinction learning is thus critical for the ability of an organism to react to environmental changes and hence to its survival (Dunsmoor et al., 2015; Goldstein and Volkow, 2011; Quirk and Mueller, 2008).

The rodent prelimbic mPFC (PL), together with the infralimbic mPFC (IL), is implicated in extinguishing reward-seeking behavior (Chen et al., 2013; Jonkman et al., 2009; Moorman and Aston-Jones, 2015; Riaz et al., 2019; Sharpe et al., 2019). Both pharmacological inactivation of PL and optogenetic stimulation of its inhibitory network during the presentation of conditioned stimuli facilitate extinction (Caballero et al., 2019; Sparta et al., 2014). Similarly, optogenetic stimulation of PL projections to the nucleus accumbens reduces reward seeking when reward is associated with risk of aversive reinforcement (Kim et al., 2017).

While such manipulations highlight the role of mPFC in extinction of reward-seeking responses, the neural dynamics driving extinction is largely unknown. At the cellular level, acquisition of new behavioral strategies is associated with changes in PL and IL activity, with the PL predicting and the IL following the acquisition of the new contingency (Rich and Shapiro, 2009). Furthermore, sudden transitions in rodent mPFC activity signal rapid behavioral shifts during rule learning (Durstewitz et al., 2010) and mark the onset of the exploratory phase following changes in cued reward probabilities (Karlsson et al., 2012). In both rodents and humans, changes in mPFC activity precede behavioral changes both for spontaneous and enforced strategy switches (Powell and Redish, 2016; Schuck et al., 2015).

Thus, while representational switches in mPFC have been studied to some degree during learning of new behavioral rules, it remains an open question whether similar dynamical processes are also at work when a previously acquired rule has to be suppressed, i.e. during extinction learning. In fact, while rule switching requires the formation of new stimulus-reward association, the loss of conditioned responses during extinction learning follows from the suppression of reward seeking *per se*. To address this question, here we analyzed *in vivo* multiple single-unit recordings from the rat PL area during maintenance and within-session extinction of a visually guided appetitive operant conditioning task. Furthermore, to enable a detailed analysis of fine-scale temporal relationships between single-unit activity, population dynamics, environmental conditions, and aspects of the animal’s behavior at a single-subject level, we combined recently developed change-point detection methods for neural activity (Toutounji and Durstewitz, 2018) with a newly developed statistical model for characterizing the temporal unfolding of transitions in the animal’s behavior. Our analyses revealed that even when experimental conditions and behavioral responses were stable, single-unit coding in the mPFC was not. Importantly, however, shortly before the animal started to actively suppress the previously acquired reward contingency, changes in single-unit activity became highly coordinated across the whole network, pushing the network toward a new internal state that drove extinction of reward-seeking behavior.

## RESULTS

### PL activity remains modulated by conditioned cues during extinction learning

We designed a visually guided appetitive operant conditioning paradigm to probe extinction of reward-seeking behavior in rats (see Methods and **Figures 1A-1B**). We chose alcohol as reward to investigate extinction learning of both appetitive reward seeking in general and drug-seeking responses in particular. Extinction therapy is, in fact, clinically used to treat substance use disorders, however with variable efficacy (Mellentin et al., 2017). To better understand the mechanisms leading to extinction of drug-seeking responses, we developed a paradigm where the amount of administered alcohol served as a positive reinforcer, shaping conditioned behavior, but did not lead to intoxication, which would have interfered with our measurements. In detail, each trial (**Figure 1B**) started with a visual cue, followed by the presentation of two levers 5 sec after cue onset, one of which, the active lever, directly below the cue light. Only responses to the active lever were reinforced by delivery of a drop of alcohol reward (40 µl, 10% v/v in water) after a 1.5 sec delay. Lever presses on the inactive lever had no consequences. The trial ended, following lever press or 10 sec after lever presentation with no response, by terminating the cue and retracting the levers. Pseudo-random inter-trial intervals (10, 15 or 20 sec) separated trials. Following habituation, appetitive conditioning, tetrode implantation (**Figure 1C**) and retraining (see Methods), a cohort of 10 rats underwent one *maintenance* session of 60 reinforced trials (**Figure 1A left**). On the next day, *within-session extinction* began with 9 reinforced trials followed by 60 unreinforced trials (**Figure 1A right**). The switch in reward contingency was not signaled to the animals and other experimental conditions were kept constant throughout the session. The number of inactive lever presses was low and comparable in both maintenance and extinction (**Figure S1A**). Response probability (rate of active lever presses) during maintenance was high, indicating that rats had learned to associate the visual cues with reward (**Figure 1D left**). This high response probability dropped during within-session extinction (60 maintenance vs. last 18 within-session extinction trials, percent of active lever presses: 87.7 ± 2.5% vs. 12.8 ± 3.1% mean ± sem, right-tailed Wilcoxon signed-rank test p = 9.8 10^−4^; **Figure 1D**), indicating that behavior was extinguished when the visual cues were not reinforced any more.

**Figure 1.**
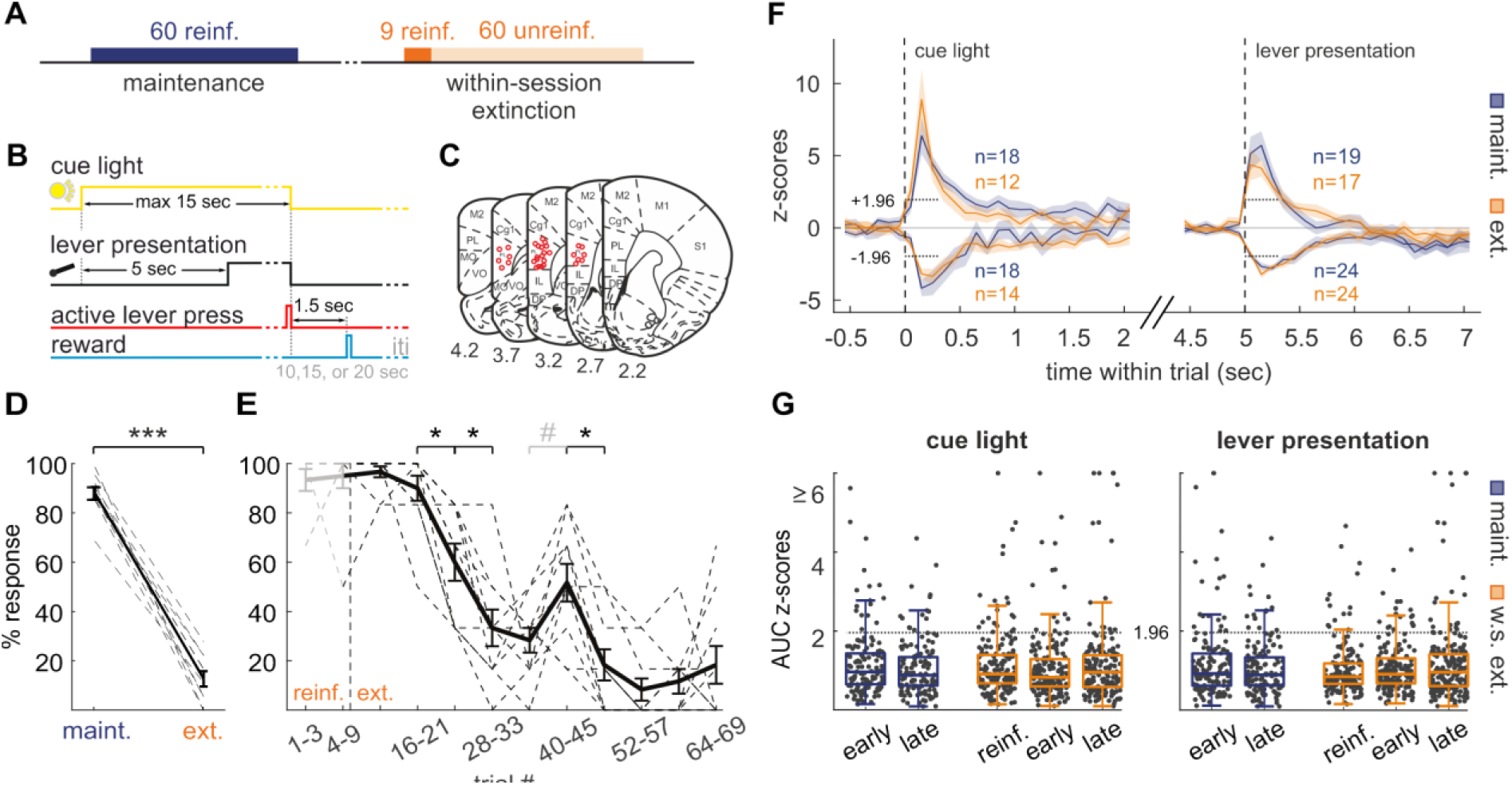
PL activity remains modulated by conditioned cues during extinction learning. (A, B) Behavioral task (A) and schematics of trial timeline during maintenance (B). During reinforced trials, reward was delivered exclusively upon pressing the cued lever (active lever). (C) Histologically verified recording sites within PL of the 10 rats. (D, E) Percentage of active lever presses during maintenance and last 18 trials of extinction (D) and throughout within-session extinction (E). Dashed lines show percentages for individual animals. Solid line and error bars show mean ± sem. Asterisk and hash symbols mark Benjamini-Hochberg corrected p < 0.05 and p < 0.08, respectively. See also **Figure S1A**. (F) z-scored activity of significantly responding units (number of units shown for each curve) following cue light and lever presentation (see Methods). Horizontal dotted lines mark the significance threshold and testing window. Solid lines and shading show mean ± sem. (G) Area-under-the-curve (AUC) for z-scored single-unit response (see Methods) computed on trial blocks of steady-state behavior (early/late: first/last 12 trials during maintenance and extinction; reinforced: 9 reinforced trials during within-session extinction). Boxplot whiskers extend to include points within 1.5 the interquartile range (IQR). Horizontal dotted lines mark the significance threshold.

In order to inspect the timing of behavioral changes during within-session extinction, responses were binned in blocks of 6 consecutive trials. We observed a gradual reduction in response probability across the whole cohort starting at trials 16-21, and an intermittent, albeit not significant, increase at trials 40-45 (Wilcoxon signed-rank tests between consecutive blocks with Benjamini-Hochberg adjusted p < 0.05 and p < 0.08, respectively; **Figure 1E**). We then investigated the PL response properties following cue and lever presentation by inspecting the z-scores of 132 and 162 units recorded during maintenance and extinction, respectively (see Methods). **Figure 1F** compares the average z-scored activity of significantly responding units during maintenance with significant z-scored activity in unreinforced trials during within-session extinction. Excitatory as well as inhibitory rate modulations following cue light and lever presentation were comparable (uncorrected Wilcoxon rank-sum test; cue light maintenance vs. extinction excitatory/inhibitory p = 0.56/0.31; same for lever presentation p = 0.50/0.48). In order to test whether the response of PL units to task stimuli is predictive of the behavioral response probability (**Figures 1D** and **1E**), we also compared activity during trial blocks of stable behavior in both maintenance and within-session extinction (first/last 12 trials during maintenance and extinction; 9 reinforced trials during within-session extinction; **Figure 1G**). We found no significant difference in z-score distributions, neither when comparing different steady-state blocks within the same session, nor when comparing corresponding blocks between the two sessions (uncorrected Wilcoxon rank-sum test; cue light maintenance vs. extinction early/late p = 0.17/0.73, early vs. late maintenance/extinction p = 0.32/0.47, reinforced vs. extinction early/late p = 0.13/0.49; same for lever presentation p = 0.68/0.53, p = 0.45/0.80, p = 0.64/0.50). These results demonstrate that, as behavior changes toward extinguishing the cue-reward association, PL remains responsive to task-related cues to a similar degree as it was when expressing the conditioned response. Moreover, during within-session extinction, the overall response to cues remains consistent across different blocks of steady-state behaviors, whether the animal responded to the task or not. These findings are in line with previous observations showing that the overall proportion of different mPFC units responding to different task aspects remains about the same despite changes in task rules and contingencies (Ma et al., 2016).

### Whole-trial PL population activity reflects behavioral changes during extinction learning

Despite demonstrating a consistent decrease in behavioral response probability across animals, the above analyses do not capture trial-by-trial changes and idiosyncrasies in each animal’s behavior, which may conceal relevant aspects of the relationship between PL activity and extinction learning. To address this, we developed a new statistical model of binary-choice behavior which captures an animal’s response-probability dynamics by a weighted sum of sigmoidal curves (see Methods). Each curve is defined by the trial at which the sigmoid is at half height (*behavioral* change point; CP_50%_), a slope that defines the rate of change per trial, and a weight specifying the amount and direction of change around behavioral CP_50%_. Statistical model selection allowed us to specify the smallest number of behavioral CPs required to explain >=95% of an animal’s behavioral variance. Such analyses revealed that the tested cohort adopted a variety of behavioral profiles that differed in the degree of abruptness of behavioral changes and in the eventual occurrence of transient reinstatements of the conditioned behavior. We found 1 or 2 behavioral CPs with descending sigmoidal curves in 6 and 3 animals, respectively, and 3 behavioral CPs with two descending and one ascending sigmoid curves in one animal (**Figures 2A** and **S1C**). We then computed the spike counts during whole trials (WT) and identified population-wide change points (population CPs) from the PL units recorded from each animal (Toutounji and Durstewitz, 2018; see Methods). Change-point detection identifies significant changes in the mean neural firing rate across trials, where the trial in which the firing rate change reached 50% of its full amplitude is identified as the *neural population* CP_50%_. In spite of the varying number of units recorded per rat, two population CPs were detected in all animals (**Figure S1C**). Visual inspection suggested that changes in response probabilities, as captured by the behavioral model, are often accompanied by population CPs. In order to quantify this relationship, we developed a statistical bootstrap procedure, based on computing a likelihood ratio statistic *λ* (see Methods). This statistic compares how strongly an animal’s change in response probability locks to its own population CPs versus an alternative set of population CPs detected in another animal. Considering all possible combinations of behavioral models and population CPs, we found a substantial bias toward positive *λ* values during extinction (*λ*_ext_; right-tailed Wilcoxon signed-rank test p = 2.0 10^−10^; **Figure 2B right**), indicating a strong match between the behavioral model and the population CPs of the same animal. As control, we also computed another set of *λ* values (*λ*_maint_) using maintenance population CPs (**Figure S1B**) and related them to the same set of extinction-behavioral models in **Figure S1C**. Contrasting the *λ*_ext_ distribution with the null *λ*_maint_ distribution (**Figure 2B**) further confirmed that behavior is significantly more locked to population CPs than expected by chance (*λ*_ext_ > *λ*_maint_; right-tailed Wilcoxon signed-rank test p = 2.0 10^−8^). Furthermore, the distribution of *λ*_ext_ for individual animals showed significantly positive values (uncorrected right-tailed Wilcoxon signed-rank test p < 0.05) in 8 out of 10 animals (**Figure 2C**). The p-values for individual animals anti-correlated with the number of recorded units (Spearman’s rank correlation coefficient p = 0.03, r = −0.68; **Figure 2C**), indicating that the non-match may be due to under-sampling PL units, rather than to differences in the neural mechanisms underlying extinction learning across animals. These analyses show that the temporal evolution of behavior during extinction learning is strongly reflected in population dynamics of PL neurons.

**Figure 2.**
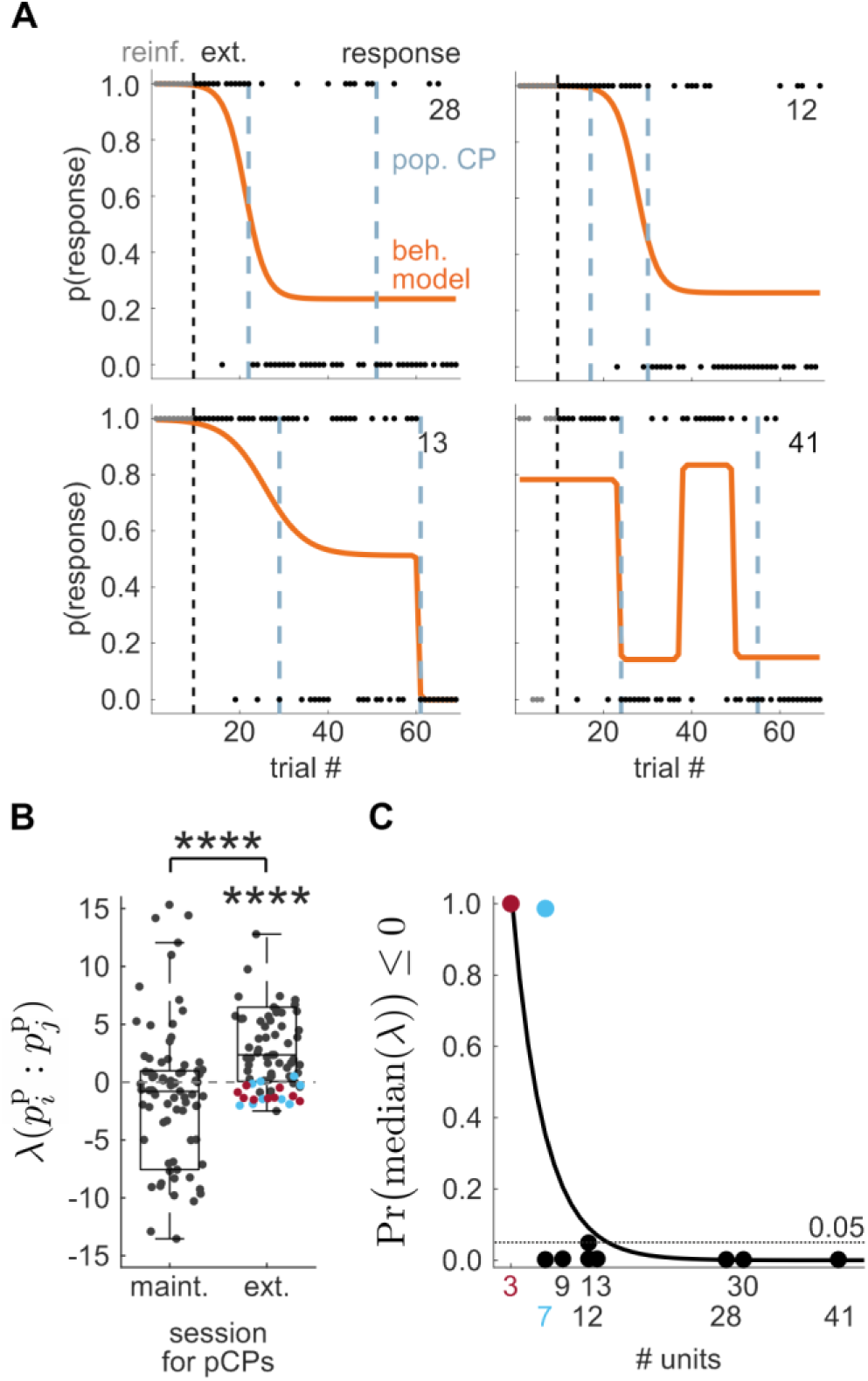
Whole-trial PL population activity reflects behavioral changes during extinction learning. (A) Examples of the behavioral models of 4 representative animals and their respective population change points (population CP) computed over the population’s whole-trial firing rate during extinction. Filled circles indicate the trial-specific behavioral choice. Dashed line indicates the onset of extinction trials. Numbers at the top-right of each panel indicate the number of recorded units. See also **Figures S1B** and **S1C** for behavioral models and population CPs of all animals during maintenance and extinction, respectively. (B) Distribution of likelihood ratio test statistic for relating the set of behavioral response models during extinction to maintenance population CPs (*λ*_maint_; left) and extinction population CPs (*λ*_ext_; right). Boxplot whiskers extend to include points within 1.5 the IQR. See also **Figures S1B** and **S1C**. (C) Number of recorded units in each rat against the p-value of its corresponding *λ*_ext_ distribution (see main text and Methods). The curve is an exponential fit that highlights the rank correlation between the two (Spearman’s rank correlation coefficient). Points in magenta and cyan pertain to *λ*_ext_ values from corresponding rats in (B) and **Figure S1C**.

### PL single-unit dynamics during extinction learning is indistinguishable from that during maintenance

While population CPs result from the activity of the whole set of recorded units and may reflect the overall dynamics of the PL network, one might expect a certain degree of heterogeneity in single-unit encoding. Indeed, change points estimated from single-unit whole-trial spike counts (*single-unit* CPs) did not always coincide with population CPs computed on the same task window (**Figure 3A**). In order to pinpoint more exactly during which task phases extinction-related changes happened and how different units were involved in them, we identified an additional set of four single-unit CPs estimated from the spike counts of four within-trial windows of interest (**Figure 3B**). The cue-light (CL) and lever-presentation (LP) windows, defined as the 0.5 sec after stimulus onset, allowed monitoring PL network responses to task-related external stimuli. The delay period (DP) window, which spans the 2 sec preceding lever-presentation, was selected to assess potential effects of reward expectation. Finally, the inter-trial interval (ITI) window, between −3 sec and −1 sec before cue-light onset, allowed considering PL dynamics independently of specific task-related activity. Importantly, none of these windows included trial periods where motor responses were expected, which allowed a fair comparison between maintenance and extinction.

**Figure 3.**
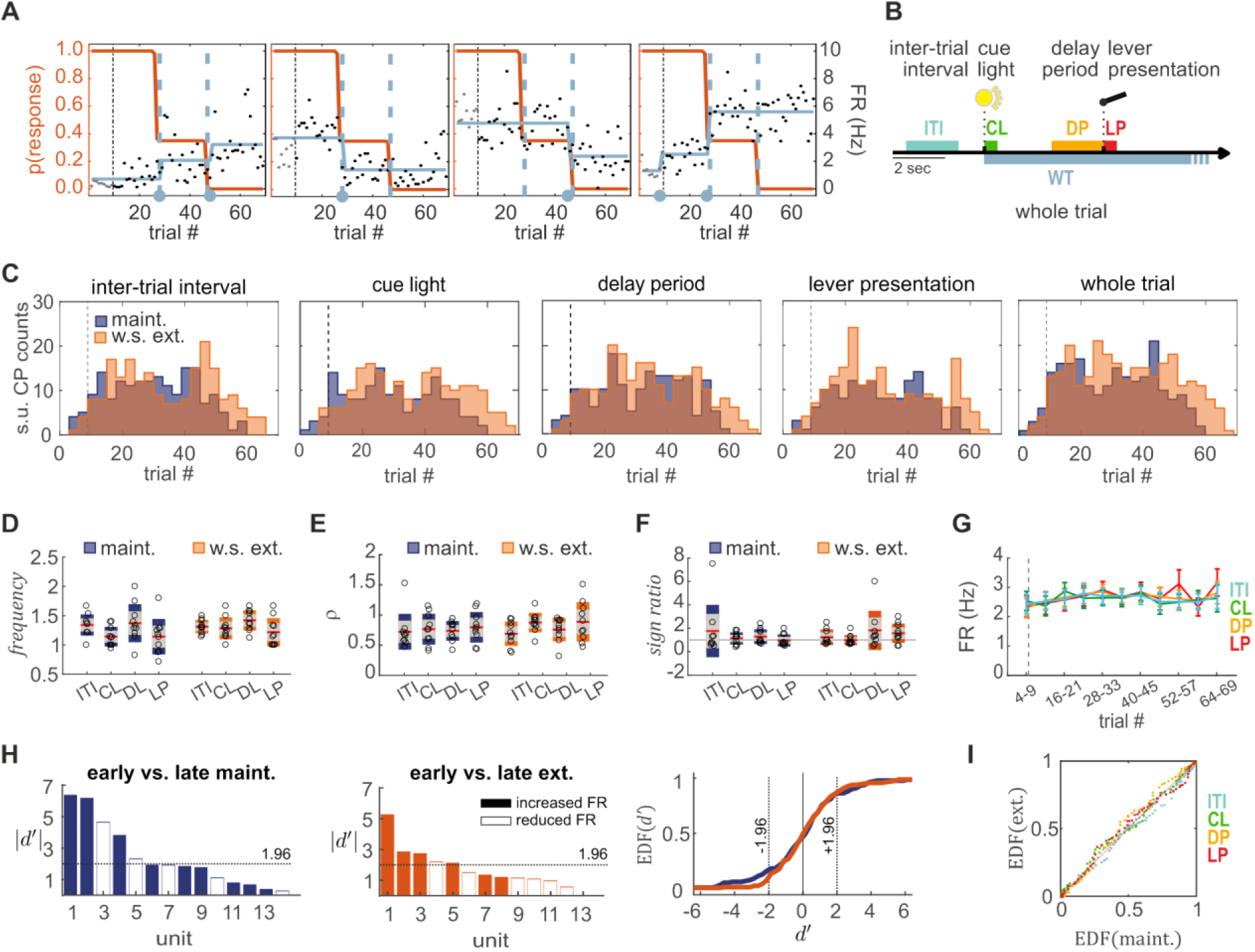
PL single-unit dynamics during extinction learning is indistinguishable from that during maintenance. (A) Four examples (same animal) of single-unit firing rates based on CP detection (gray solid line) and corresponding single-unit CPs (gray filled circles), identified from whole-trial firing rate (black dots). Behavioral model and whole-trial population CPs are as in **Figure 2A** and **S1C**. (B) Five sets of single-unit CPs identified from single-unit firing rates within five task windows defined relative to light onset as: inter-trial interval (ITI; sec −3 to −1), cue light (CL; sec 0 to 0.5), delay period (DP; sec 3 to 5), lever presentation (LP; sec 5 to 5.5) and whole trial (WT; sec 0 to 15). (C) Distribution of single-unit CPs across maintenance and extinction trials (60 and 69 trials, respectively) for each task window, pooled from all animals. (D-F) Number of single-unit CPs per unit (D), relative change in firing rate (E), and positive-to-negative sign ratio (F), computed in four task windows. Plots show mean ± 1.96 · sem (red and gray) and sd (blue or orange). Open circles indicate the mean for individual animals. The three quantities (D-F) are statistically indistinguishable when compared between sessions. Gray line in (F) marks a sign ratio of 1, where positive and negative rate changes are balanced. (G) Population firing rate per task window over blocks of 6 consecutive extinction trials (trials 1 to 3 excluded; cf. **Figure 1E**). Dashed line indicates the onset of extinction trials. Solid lines and error bars show mean ± sem. (H) Sensitivity analysis showing increased and decreased single-unit whole-trial firing rates of a representative animal during the first and last 12 trials of maintenance (left) and extinction (middle). Empirical distribution functions (right) of the sensitivity index *d*′ for all recorded single-units in maintenance (blue) and extinction (orange) show no significant difference, despite difference in behavior. Dotted lines mark the threshold of significant change in firing rate. See also **Figure S2.** (I) P-P plot comparing the empirical distribution function of single-unit CPs over maintenance and within-session extinction trials (cf. (C)) for the four task windows.

Irrespective of the window examined, single-unit CPs during extinction learning were distributed across the entire session (ITI/CL/DP/LP/WT; **Figure 3C**). On average over all recorded units, single-unit CPs occurred in all windows with similar *frequency*, same relative change in firing rate *ρ*, and with balanced positive to negative *sign ratio* (see Methods for formal definitions of these three statistics; *frequency*: one-way ANOVA on task windows, main effect F(3,36) = 2.0 p = 0.1; **Figure 3D right**; *ρ*: one-way ANOVA on task windows, main effect F(3,36) = 1.7 p = 0.2; **Figure 3E right**; *sign ratio*: t-test on the fraction of positive over negative jumps against 1, uncorrected p > 0.05 for all windows; **Figure 3F right**). Besides, the average PL firing rate remained constant throughout the whole session for all task windows (see Methods; one-way ANOVA for repeated measures to test the effect of the trial block on the population firing rate: ITI: F(10,90) = 0.6 p = 0.8; CL: F(10,90) = 0.7 p = 0.7; DP: F(10,90) = 0.9 p = 0.5; LP: F(10,90) = 1.9 p = 0.1; **Figure 3G**). While overall changes in firing rate across units were balanced, the specific changes in single-unit firing rates resulted in a reorganization of the PL coding throughout the session: While the unit responses within CL and LP phases remained unchanged on average (across the population) from the beginning to the end of the extinction session (cf. **Figure 1G**), the identity of task-responsive units varied. About 21.6% of the recorded units changed the trial-averaged firing rate by more than 2 standard deviations between the first and last 12 extinction trials of the session (sensitivity index *d*′, see Methods; **Figures 3H middle** and **S2B**). Only 30% of the units with significant response in the first reinforced trials of within-session extinction (40% and 20%, following CL and LP, respectively) were also responsive at the end of the session (cf. **Figure 1G**). Surprisingly, this degree of change in task-responsive units was also present during maintenance, where both experimental conditions and animal behavior were constant throughout the session (**Figures 3D-3F left** and **3H left**): We found no significant difference between maintenance and extinction single-unit CPs for all windows (ITI/CL/DP/LP) with regard to the distribution of CP *frequency, ρ, sign ratio*, and the distribution of CP occurrence across the trials of each session (*frequency*: repeated measures two-way ANOVA, factors: session/window, main effects: session F(1,9) = 1.4 p = 0.3, window F(3,27) = 5.9 p = 0.003, interaction F(3,27) = 0.5 p = 0.7; **Figure 3D**; *ρ*: repeated measures two-way ANOVA, factors: session/window, main effects: session F(1,9) = 0.5 p = 0.5, window F(3,27) = 1.6 p = 0.2, interaction F(3,27) = 0.4 p = 0.7; **Figure 3E**; *sign ratio*: two-way factors: session/window, main effects: session F(1,8) = 0.13 p = 0.7, window F(3,24) = 1.6 p = 0.2, interaction F(3,24) = 1.5 p = 0.2; **Figure 3F**; post hoc Bonferroni correction applied in all ANOVAs, no significance found post hoc; single-unit CP trial distribution: see Methods, 2-sample Kolmogorov-Smirnov test for single-unit CPs in ITI: p = 0.4; CL: p = 0.4; DP: p = 0.1; LP: p = 0.4; **Figure 3I**, cf. **Figure 3C**). Moreover, similar to extinction, a large fraction of units changed their trial-averaged firing rate from the start to the end of the maintenance session (30.3%; **Figures 3H left** and **S2A**). The distributions of the sensitivity index *d*′ for units recorded in the maintenance versus extinction sessions were, in fact, not significantly different (2-sample Kolmogorov-Smirnov test p = 0.5; **Figure 3H right**).

In summary, PL units changed their responsiveness to the task stimuli with balanced positive and negative single-unit CPs during both maintenance and extinction. This reorganization occurred to the same extent in both these sessions and therefore was not induced by changes in the experimental conditions.

### PL baseline rate and task-evoked responses change in anticipation of behavioral extinction

The above analysis revealed that, during prolonged stretches of time (every session lasted about 30 min), PL single-units changed their responsiveness to stimuli. Such variation in firing patterns also occurred during maintenance and the number of single-unit CPs was comparable in the two sessions. Therefore, we wondered whether the observed match between population and behavioral CPs during extinction was due to a coordination of single-unit CP occurrences that was locked to the change in behavior, rather than to an overall increase or decrease in firing rates. We found that, despite the similarity in single-unit CP-related statistics between the two sessions, at the population level the number of population CPs per rat was significantly higher during extinction, irrespective of the task window considered (two-way ANOVA: main effect of session: F(1,72) = 67.8 p = 5.7 10^−12^, main effect of task window: F(3,72) = 7.0 p = 0.6. Interaction F(3,72) = 0.7 p = 0.6; **Figure 4A**).

**Figure 4.**
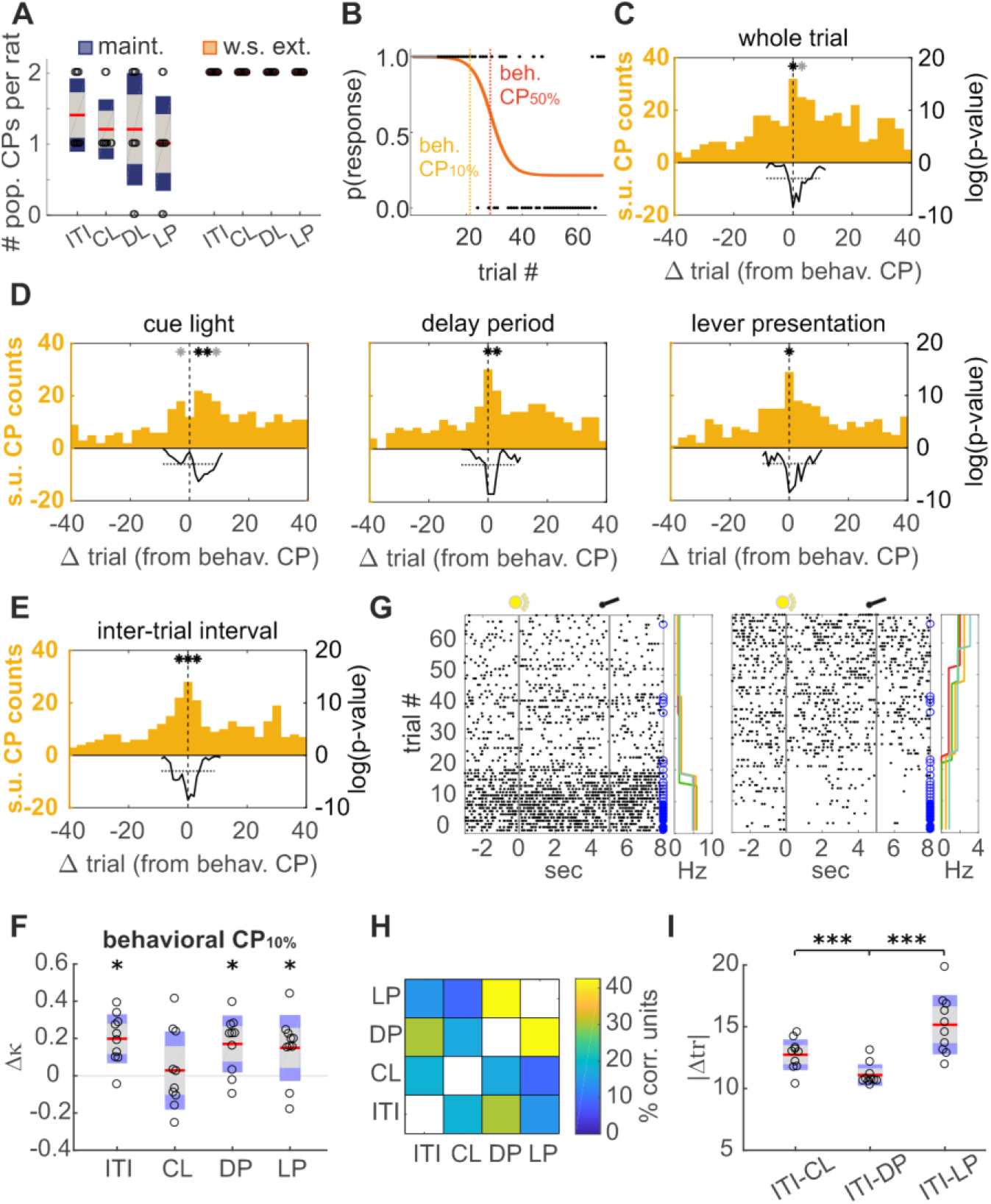
PL baseline rate and task-evoked responses change in anticipation of behavioral extinction. (A) Number of population CPs per animal. Plots show mean ± 1.96 · sem (red and gray) and sd (blue or orange). Open circles indicate the numbers for individual animals. (B) Onset (yellow) and center (red) of an extinction episode for one representative animal. Behavioral CP_10%_ and behavioral CP_50%_ correspond to 10% and 50% drop in response probability, respectively. Behavioral choice and model are as in **Figure 2A** and **S1C**. Also see **Figure S4A**. (C-E) Single-unit CP distributions for different task windows (WT/CL/DP/LP/ITI) pooled across animals and aligned with respect to each animal’s behavioral CP_10%_. Single-unit CPs coordinated at extinction onset in all windows. Statistical test performed via bootstrap (see Methods). Asterisks mark Benjamini-Hochberg corrected p-values with p < 0.05 (black) and p < 0.1 (gray). Black solid lines trace the log(p-value). Horizontal dotted lines mark the log(0.05) threshold over the tested window. Also see **Figure S3**. (F) Classifier performance in predicting the animal’s behavioral state from population firing rates during four task windows (cf. **Figure 3B**). Significance was assessed via bootstrap (see Methods). Differences between data and bootstrapped Cohen’s kappa are reported by showing mean ± 1.96 · sem (red and gray) and sd (purple). Open circles indicate differences for individual animals. Population rates are predictive of extinction onset in ITI, DP and LP. See **Figure S4C** for a similar analysis using behavioral CP_50%_. (G) Raster plots of two representative units from the same animal, showing rate progression across extinction (bottom to top). Filled and open blue circles mark trials with reinforced and unreinforced lever presses, respectively. Vertical gray lines indicate cue light onset and lever presentation. To the right, single-unit firing rates based on CP detection (cf. **Figure 3A**) in four task windows, color-coded as in **Figure 3B**. (H) Fraction of single-units for which the evolution of firing rates within one window significantly correlates with that within a second window. (I) Firing-rate changes during ITI are most coordinated with those occurring during DP. Absolute distance in trials between the occurrence of a single-unit CP in ITI and the closest single-unit CP of the same unit in CL, DP and LP. Plots show mean ± 1.96 · sem (red and gray) and sd (purple). Open circles mark values for individual animals.

To test the hypothesis that single-unit CPs are coordinated with behavioral change, we considered for each animal the lag between its single-unit CPs and the onset of an extinction-learning episode. Extinction onset was defined as the trial correspondent to a decrease by 10% (behavioral CP_10%_) in response probability as captured by the behavioral models (see Methods and **Figure 4B**). We found that single-unit CPs computed over the whole trial locked with zero lag to behavioral CPs_10%_ (see Methods and **Figure 4C**). A similar match was also found for single-unit CPs occurring during the CL, DP and LP windows (**Figure 4D**). To further confirm the temporal coordination between single-unit CPs and the learned behavior, we performed, as a control, the same analysis but matching behavioral CPs of extinction to the single-unit CPs of maintenance instead. Besides an increase in single-unit CP occurrence probability at the center of the maintenance session, which explains the non-uniform distribution of maintenance single-unit CPs around behavioral CPs, no fine-tuned coordination was found (Benjamini-Hochberg corrected bootstrap test; see Methods and **Figure S3A**). Behavioral extinction thus coincided with a coordinated change in PL-unit firing rates. Such coordination preceded the change in behavior. In fact, while the behavioral CP_10%_ indicates a decrease in response *probability* from baseline of 10%, 7 out of 10 animals responded to all lever presentations until, and including, the first behavioral CP_10%_ mark (**Figures S4A** and **S4B**). As a further confirmation, we found in all four task windows considered that the single-unit CP probability significantly increased a few trials prior to behavioral CP_50%_ (**Figure S3B right**).

Interestingly, also during the ITI, when the animal was not actively engaged in the task, PL firing rate changed in anticipation of behavioral CPs (**Figure 4E**). To further confirm whether behavioral changes could be predicted from the activity of PL neurons even before the beginning of a trial, we trained a classifier on PL population spike counts during the ITI window preceding trial t to predict whether the animal will still be committed (t < behavioral CP) or not (t > behavioral CP) to the task on that trial. We measured the classifier performance using Cohen’s kappa (where *κ* = 1 corresponds to a perfect prediction, *κ* = 0 to chance and *κ* = −1 to complete mismatch; see Methods). We found that, based on population vectors constructed from the ITI, *κ* = 0.45 ± 0.08 when considering behavioral CP_50%_ and *κ* = 0.49 ± 0.07 when considering behavioral CP_10%_. To confirm that the observed effect was due to behavioral extinction and not, more generally, to random monotonic changes in firing rate across the session, we constructed bootstrap replica where unaltered PL population vectors were used to predict the occurrence of randomly generated behavioral CPs (see Methods). PL activity during the ITI could predict the original behavioral CPs significantly better than the bootstrapped replicas (one-tailed paired t-test, p = 5.2 10^−4^; **Figure 4F**). A similar result was obtained when considering population spike counts during DP and LP (one-tailed paired t-test, Benjamini-Hochberg adjusted over the four task windows, p ≤ 0.012 for all task windows except CL; **Figures 4F** and **S4C**). PL population activity during CL was at chance level in predicting behavioral CP_10%_ (**Figure 4F**), but above chance when predicting behavioral CP_50%_ (one-tailed paired t-test, Benjamini-Hochberg adjusted over the four task windows, p ≤ 0.01 for all task windows; **Figure S4C**). This was in line with the observed lags in temporal coordination between single-unit CPs during CL and either behavioral CP_10%_ or behavioral CP_50%_ (cf. **Figures 4D left** and **S3B right**, respectively), suggesting that changes in PL response to cue light occurred between a drop of 10% and 50% in an animal’s response probability.

The previous analyses showed that behavioral CPs were anticipated by a coordinated change in firing rate that affected all task windows. Moreover, the ability to predict behavior from population vectors did not differ between task windows (repeated measures one-way ANOVA on Δ*κ* computed on the four windows: main effect F(3,27) = 1.3 p = 0.3; cf. **Figure S4C**). These results suggest that behavioral extinction corresponds to a global reorganization of network activity across all task phases, rather than to a modulation of unit responses to specific conditioned stimuli. **Figure 4G** shows raster plots of two exemplary units from the same animal, depicting how changes in baseline firing rate across the whole trial occurred in coordination with behavioral extinction, both in stimulus-responsive (**Figure 4G right**) and non-responsive (**Figure 4G left**) units. Firing-rate changes during the four task windows of interest (analyzed pairwise) correlated in about 22% of the recorded units (Spearman’s correlation, Benjamini-Hochberg adjusted p < 0.05; **Figure 4H**). Notably, despite the ITI window being positioned between the LP and CL windows of two successive trials, the firing rate during the ITI was significantly correlated for more units (30%) with that during DP than that during the CL (19%) and LP (14%) phases. Since changes in firing rate were better coordinated between the ITI and DP than between the ITI and any other task phase, we expect that single-unit CP occurrence was comparably better aligned between the ITI and DP windows than between the ITI and other task phases. To evaluate this hypothesis, we tested whether the lag between the occurrence of single-unit CPs during the ITI and single-unit CPs during other task windows was comparable for the three windows (see Methods). As expected from **Figure 4H**, single-unit CPs during the ITI were most coordinated with those during DP (repeated measures one-way ANOVA with Greenhouse-Geisser correction: main effect F(1.1,10.3) = 17.2 p = 0.001, post hoc with Bonferroni correction: (ITI-CL, ITI-DP) p = 0.0008, (ITI-LP, ITI-DP) p = 0.001, (ITI-CL,ITI-LP) p = 0.07; **Figure 4I**). This agrees with previous observations that rate changes in accordance with reward expectations were particularly prominent within delay phases, during which the animal neither had to process specific sensory stimuli nor to initiate specific responses (Leon and Shadlen, 1999; Watanabe, 1996). In our extinction paradigm, conditioned stimuli were identical for each trial, with only the active lever being cued throughout the session. Thus, it is possible that animals familiar with the task encoded reward expectation in PL neurons, not only during DP, but also during ITI.

In summary, PL encoded strategy changes by coordinating single-unit CPs in anticipation of behavioral changes. Single-unit CPs mark longer-term transitions in a unit’s firing rate, and hence the population-wide coordination of such events indicate a complete reorganization of the neural activity pattern as indicative of a transition between attractor states. Consistent with this idea, rate changes were not limited to any particular task phase, or even to the proper trial periods, but were consistent across all phases including the ITI, and hence marked a global transition in the prefrontal ensemble state, possibly driven by updates in reward expectancy.

## DISCUSSION

Lesion (Fragale et al., 2016), pharmacological inactivation (Caballero et al., 2019; Ramanathan et al., 2018) and optogenetic stimulation (Marek et al., 2018; Sparta et al., 2014) studies provide increasing evidence that PL is a critical locus for extinction of reward-seeking behaviors and conditioned fear. Using an appetitive operant within-session extinction paradigm, we studied the dynamics of PL leading to the extinction of conditioned reward-seeking behavior in rats. By using newly developed process model and statistical tools for change detection to examine idiosyncrasies in each animal’s behavior and neuronal activity, we were able to identify and parametrize trial-to-trial changes in both. This revealed that fine temporal coordination in mPFC dynamics guides extinction learning.

During across-session extinction, mPFC responsiveness to conditioned stimuli persists over subsequent unreinforced days (Moorman and Aston-Jones, 2015). We showed here that also during within-session extinction, the population response to conditioned task stimuli remained equally strong even when the animal stopped acting upon them. This sustained responsiveness did not correspond, however, to stability in single-unit coding. While, on the one hand, some single units in mPFC can maintain their response pattern across days (Brebner et al., 2020; Powell and Redish, 2014), on the other hand, we found that the majority of units significantly changed their average firing rate and their stimulus-responsiveness within tens of minutes. It is important to note that our approach captures longer-lasting changes in unit coding, i.e., significant jumps in firing probability between periods of relative stability, and not trial-to-trial variability as observed in many brain regions. Changes in unit coding were widespread along the session and occurred also under stable experimental conditions and stable behavioral responses. In fact, changes in single-unit coding do not necessarily imply changes in population coding, where coding properties might be preserved by redundancies in the ensemble representation (Hirokawa et al., 2019; Narayanan et al., 2005; Puchalla et al., 2005) or within consistent neural trajectories (Enel et al., 2016; Mante et al., 2013). The fact that we observed the same degree of single-unit changes during maintenance and within-session extinction, which imply different cognitive demands, may therefore suggest that these transient representations are an intrinsic feature of prefrontal dynamics, rather than the result of specific cognitive demands.

Upon first inspection, we found no discernible difference between PL activity during maintenance and within-session extinction, specifically in regards to the rate and likelihood of single-unit facilitation and suppression. A more detailed analysis revealed, however, a strong temporal coordination across the entire population during extinction learning, which was not present when no learning was required of the animal during maintenance. These two effects, the reorganization of PL activity irrespective of learning and the population-level temporal coordination specific to the learning phase, resonate with recent findings on PL plasticity during sleep in response to rule learning (Singh et al., 2019), suggesting that similar neural mechanisms may underlie both the formation and fading of response-reward associations in PL. Our finding that coordinated PL reorganization anticipates change in behavior toward extinction lends further credence to this hypothesis. In fact, ample evidence shows that in rat prefrontal cortex, particularly its prelimbic subregion, neural population dynamics is reshaped prior to rule and reversal learning of appetitive reward-seeking strategies (Durstewitz et al., 2010; Karlsson et al., 2012; Powell and Redish, 2016; Rich and Shapiro, 2009). In light of those studies, our results may suggest a causal link between PL coordination and strategy switching, irrespective of the particular mode of learning, i.e. whether a new rule is acquired or an old one needs to be suppressed.

Network reorganization in PL during extinction learning was a global property not anchored to a particular cognitive phase of the task. Temporally coordinated changes in neural activity were observed during different trial stages, including resting periods between trials, rather than being confined to specific windows within the trial. In fact, irrespective of the task window considered, population firing rates predicted changes in animal behavior equally well. In the presence of ambiguous sensory information, successful action selection is based on forming a reliable model of the environment as represented by the animal’s *belief states* (Babayan et al., 2018). Lesion studies have shown that mPFC plays a fundamental role in the computation of belief states and in cognitive control (Gershman and Uchida, 2019; Ridderinkhof et al., 2004; Sharpe et al., 2019). Our results suggest that updating belief regarding the availability of reward following extinction may correspond to a shift within phase space of the network’s resting state. Theoretical and experimental works support the presence of attractor dynamics in PFC (Durstewitz et al., 2010; Katori et al., 2011; Wimmer et al., 2014). Specifically, Redish and colleagues pushed forward the hypothesis that the prolonged absence of an expected reward would lead to the formation of a new attractor state in the mPFC representing the changed contingencies (Redish et al., 2007). Within this framework, a shift in phase space as suggested by our data may correspond either to a transition between two pre-existing attractor states, led by external inputs, e.g., from the hippocampus (Sotres-Bayon et al., 2012) or the amygdala (McGinty and Grace, 2008; Senn et al., 2014), or to the formation of a new attractor state through plasticity (Dunsmoor et al., 2015; Toutounji and Pipa, 2014) or neuromodulatory processes (Harris and Thiele, 2011). Upon inspecting single-unit spike trains during extinction trials, we observed units within the same network with both slow and abrupt rate changes. This may suggest a third possible scenario where learning slowly modulates the activity of a few neuronal assemblies, possibly upon updates of reward expectancy, which lead network dynamics to undergo an abrupt transition between two global attractors as the more slowly changing units accumulate evidence for different contingency scenarios.

Beyond providing insights into motivational processes and learning, understanding reward extinction-learning mechanisms also carry translational value for addiction research. Hence, while seeking reward is fundamental for survival, excessive drug-seeking following cue exposure is a central component of addictive behavior. One behavioristic psychological approach to treat alcoholics or drug addicts is cue exposure therapy (CET - i.e., extinction therapy). In CET, patients are exposed to relevant drug cues to extinguish conditioned responses. CET shows varying degrees of efficacy (Mellentin et al., 2017) and therefore it is of critical importance to understand its underlying neurobiological mechanisms. Our results indicate that extinction of alcohol-seeking behavior is not associated with a loss in mPFC responsiveness to conditioned stimuli. Instead, extinction manifests as a network-wide transition between two states corresponding to distinct behaviors: response (or consumption) and omission (or abstinence). This observation may thus suggest an alternative approach towards a pharmacologically driven CET that targets the relative strength and stability of the neural attractors representing the consumption and omission states. Such an intervention may go in two directions, either weakening the consumption state to facilitate extinction, thus avoiding maladaptive persistence in harmful behaviors, or strengthening the abstinence state to reduce the triggering effect of conditioned stimuli, thus reducing their valence and attenuating their ability to induce relapse.

## METHODS

### Animals

Two-month-old male Wistar rats (Charles River, Germany) were group-housed in standard rat cages under a 12h/12h reversed light/dark cycle. Food and tap water were provided *ad libitum*. After tetrode implantation, rats shared a cage in groups of two, separated by a high, perforated wall (50 cm) allowing snout contact. All experimental procedures were performed in accordance with the EU guidelines for care and use of laboratory animals and were approved by the local Committee (G-273/12 and G30/15; Regierungspräsidium Karlsruhe, Germany).

### Behavioral training

All self-administration training sessions were carried out 2 h after the beginning of the dark phase in operant chambers (ENV-008CT; interior: L, 30.5 cm; W, 24.1 cm; H, 21.0 cm; Med Associates Inc.; VT, USA). These training chambers were located inside sound-attenuating cubicles containing a white-noise generating fan (ENV-025F28). Training consisted of 3 steps. In step 1 only one lever was presented, and the rats underwent 4-5 sessions of behavioral shaping (water-deprived for 20 h before the first two sessions) until they reached a maximum of 50 drops (30 μl per drop) of 10% alcohol (v/v in water) under a fixed ratio 1 schedule in maximally 1 h. In step 2, the rats were trained to self-administer 10% alcohol in sessions with 60 trials on a fixed inter-trial interval (15 s) schedule for five days. In these sessions, the lever was presented for maximally 10 s and was retracted immediately following lever press. In step 3, a cue light above the active lever, an inactive lever on the opposite side and a variable inter-trial interval (10, 15 and 20 s) were introduced. The cue light was first presented for maximally 15 s and, 5 s after cue light onset, levers were presented for maximally 10 s. Response on either lever terminated the light and caused levers to retract, but only a response to the active lever was deemed successful and followed by delivery of alcohol. These training sessions consisted of 60 trials each and lasted for 3-4 weeks. Performers with success rates > 50% were selected to continue training in the intended recording chamber for about another week. Rats with stable performance >70% on two subsequent days were selected for tetrode implantation (see Surgery). The recording chamber (ENV-007CT; interior: L, 30.5 cm; W, 24.1 cm; H, 29.2 cm) was higher than the training chamber to make room for an electrical swivel commutator (Dragonfly Inc., USA), allowing data acquisition in freely moving animals. In this chamber, one drop contained 40 μl 10% alcohol and was supplied in a cup by a motor-driven liquid dipper (ENV-202M-S), causing a delay of 1.5 s after active lever press. Head entries through the liquid dipper’s access opening (5.08 x 5.08 cm) were detected by interrupting an infrared beam across entrance (Med Associates Inc.; VT, USA), and the behavior was observed with a USB camera (Delock 95353; Conrad, Germany).

### Surgical and tetrode placement procedures

Rats were anesthetized with isoflurane (1.5-2.0%). Custom-built flexDrive (Voigts et al., 2013) containing 8 tetrodes (12.5 µm Teflon-coated tungsten wire, California Fine Wire) was unilaterally implanted with a 10-degree angle towards midline into the prelimbic part of the mPFC (PL: A/P, +2.8 to +3.8 mm; M/L, +0.8 to +1.3 mm; D/V, −2.5 to −2.6 mm). A bone screw above the cerebellum served as ground. The craniotomy was stepwise sealed with three-component adhesive (Super-Bond C&B; MPE Dental UG, Wesseling, Germany) and two-component embedding resin (Technovit 5071; Kulzer Wehrheim, Germany). From the next day after surgery, tetrodes were advanced gradually every second day. The location of the tetrodes within PL was confirmed (**Figure 1C**) in fixed, 50 µm thick, coronal sections by Nissl staining following current passing (100 µA; 20 s) to deposit iron particles via a Prussian blue reaction (Ma et al., 2016).

### Behavioral task

Rats with single-unit activity were retrained in the recording chamber with 60 reinforced trials for 3-5 days until reaching > 75% average success rates on three consecutive days (n=10; 86.3 ± 1.7% mean ± sem). On the next day, within-session extinction began with 9 reinforced trials followed by 60 unreinforced trials.

### Recording

Multiple single-units were simultaneously recorded using a 32-channel RHD2132 amplifier connected to a RHD2000 USB interface board (Intan Technologies LLC, CA, USA). All channels were digitized with 16-bit resolution, sampled at 30 kHz and band-pass filtered between 0.1 Hz and 8000 Hz. The time stamps for external stimuli (cue light, lever presentation), lever presses, dipper activation, and head entries into the liquid dipper’s access opening were transmitted from the Med Associates behavioral control system (Med-PC IV software, version 4.39; Med Associates Inc.; VT, USA) to the Intan recording system to align behavior to neural activity.

### Spike detection and sorting

After band-pass filtering between 300 and 5000 Hz (4th order Butterworth filter, built-in MATLAB function), the median voltage trace of all channels was subtracted from each trace to reduce noise. Out of the three consecutive days with > 75 % mean success rate during retraining, the day with the highest number of single-units was chosen for further analyses (maintenance). The threshold for spike detection was set at 5.5 times the median absolute deviation from baseline. Detected spikes were sorted with a custom-built graphical user interface in MATLAB (provided by W. Kelsch, CIMH Mannheim) into individual cell clusters based on peak amplitude and the first three principal components of the waveform. Spike sorting quality and unit isolation were assessed with MLIB, a MATLAB (Mathworks) toolbox for analyzing spike data by Maik Stüttgen (https://www.mathworks.com/matlabcentral/fileexchange/37339-mlib-toolbox-for-analyzing-spike-data). After spike sorting, less than 1% of consecutive spikes in accepted clusters had an interspike interval < 2 ms. Cross-correlation analyses supported that each single-unit was isolated from other units.

### Data analysis

All data were analyzed using built-in and custom-made MATLAB routines (Mathworks), so was model fitting. To correct for multiple comparisons, significance levels were adjusted using the Benjamini-Hochberg procedure (Benjamini and Hochberg, 1995). ANOVAs were performed using the SPSS software (IBM).

### Z-scores

We obtained single-unit instantaneous firing rate as follows: Spike trains were first convolved with a Gaussian kernel (s.d. = 60 ms) and the resulting time series were time-averaged within 100 ms bins. Binning was aligned to the cue onset of each trial. The z-scored response over the relevant block of trials (60 trials in **Figure 1F** and 12 or 9 trials in **Figure 1G**; see Results for details) was then computed for each unit by first trial-averaging (mean FR across a block of trials), followed by subtracting the mean and dividing by the standard deviation of baseline trial-averaged firing rate (2 seconds prior to either CL or LP). Significant units in **Figure 1F** were identified as those whose absolute average z-scored response over 3 bins following the stimulus (CL or LP) is above 1.96, i.e., a value outside the 95% confidence interval of the standardized normal distribution. In **Figure 1G**, the area-under-the-curve (AUC) of the z-scored response is similarly computed as the absolute average over 3 bins.

### Behavioral change model

We formally treat the behavior of each animal as a binary vector **y** = *y*_1_,…, *y*_*T*_, with *T* the number of trials in a session (*y*_*t*_ = 1 for active lever press and *y*_*t*_ = 0 for omission or inactive lever press). We thus use an inhomogeneous Bernoulli process **x** = *x*_1_,…, *x*_*T*_ to model response probability,

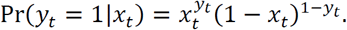

The time-varying response probability *x*_*t*_ ∈ [0,1] is, in turn, modelled as a weighted sum of *B* logistic sigmoids,

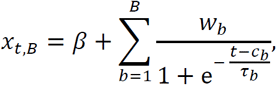

where *β* is baseline response probability. Each sigmoid is parametrized by a weight *w*_*b*_, a center *c*_*b*_ and a time constant *τ*_*b*_. Centers *c*_*b*_ correspond to *B* behavioral change points (behavioral CP_50%_’s). Model parameters are initialized using the PARCS method for any given *B* (Toutounji and Durstewitz, 2018), then inferred by constrained maximization of data log-likelihood,

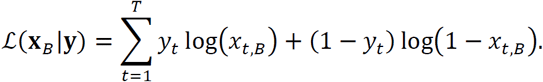

Baseline and weight constraints are imposed to assure that response probability is bounded between 0 and 1. Other parameter constraints (1 ≤ *c*_*b*_ ≤ *T* and *τ*_*b*_ > 0) enforce model identifiability. Model selection relies on an iterative procedure where, starting from *B* = 0, a null model of order *B* is compared against the order *B* + 1 alternative, using the following likelihood-ratio test,

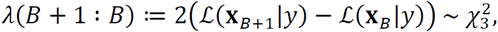

where significance level is set to *α* = 0.05. The number of degrees of freedom corresponds to the difference in number of parameters between the two models. In **Figures 2A** and **S1B** and **S1C**, we report the best models, optimized given each animal’s behavior.

### Change-point analysis

Both the initialization of center parameters of the behavioral models before optimization and neural CP (population CP/single-unit CP) detection were performed using PARCS (Toutounji and Durstewitz, 2018; **Figures 2A, 3A, S1B** and **S1C**). For neural CPs *c*_*n*_ the method is applied to the square-root-transformed spike counts in each window of interest, bringing counts closer to a Gaussian distribution and stabilizing the variance (Kihlberg et al., 1972). An upper bound of 3 on the number *N* of CPs per unit or population is chosen and the nominal significance level for the method’s permutation bootstrap procedure is set to *α* = 0.2 in order to correct for the method’s conservativeness (Toutounji and Durstewitz, 2018).

### Relating behavioral model to population CPs

In order to quantify locking between population CPs and behavior, we developed a measure for comparing the likelihoods that two sets of population CPs are sampled from one behavioral response probability distribution *p*^b^(*y*_*t*_ = 1) (**Figures 2B** and **2C**). This distribution is the sum of *B* bell-shaped curves (each peaking at one behavioral CP_50%_ *c*_*b*_ and of width that scales with *τ*_*b*_), computed by normalizing the first differences |Pr(*y*_*t*+1_ = 1|*x*_*t*+1,*b*_) − Pr(*y*_*t*_ = 1|*x*_*t,b*_)| to sum up to 1. Similarly, a neural response probability distribution *p*^p^(ΔFR_*t*_ ≠ 0) is computed as the sum of *N* Dirac delta functions, centered at the *N* population CPs *c*_*n*_. Weights are computed by averaging and normalizing 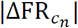 over the whole population, such that *p*^p^(ΔFR_*t*_ ≠ 0) sums up to 1. Given two neural response distributions 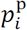 and 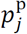, we compute the likelihood ratio,

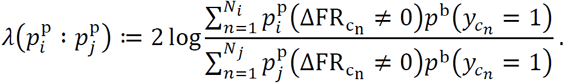

Positive 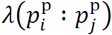 indicates a stronger locking to the same behavior of the set *i* of population CPs, relative to the set *j*. In **Figure 2B right**, we compute 9 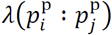 for each of the 10 animals, where *p*^b^ and 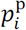 correspond to the animal’s own response probability and neural response distributions of the extinction session, respectively, and 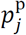 to the neural response distribution of each of the other 9 animals (*i* ≠ *j*). The resulting 10 sets of 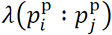 values are used to test whether *λ* > 0 for each individual animal (uncorrected right-tailed Wilcoxon signed-rank test, p-values in **Figure 2C**). To generate the null distribution of 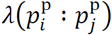 In **Figure 2B left**, we use the neural response distributions of the maintenance session and response probability distributions of the extinction session.

### Characteristics for comparing different CP sets

We compare different sets of single-unit CPs using the statistics: *frequency*, # s. u. CP/# units; relative rate change, *ρ* := 2|FR_pre s.u.CP_ − FR_post s.u.CP_|/(FR_pre s.u.CP_ + FR_post s.u.CP_); *sign ratio*, # positive s. u. CPs/# negative s. u. CPs. The firing rates FR_pre s.u.CP_ and FR_post s.u.CP_ were computed over the periods of constant firing rates around the single-unit CP, defined by PARCS as the longest periods before and after a CP where no other change point was detected. Statistics for the maintenance and extinction sessions are reported in **Figures 3D** and **3F**.

### Sensitivity analysis

We computed for each unit the statistic,

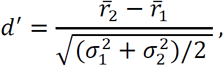

where 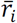 and 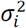 are the firing rate mean and variance over a block of trials, respectively (first vs. last 12 maintenance and extinction trials). Positive *d*′ indicates an increase of firing rate in block 2 compared to block 1 measured in units of average standard deviation, and vice versa (**Figure 3H** and **S2)**.

### Comparing single-unit CP distributions across sessions

The P-P plots of **Figure 3I** compare the empirical distributions function (EDF) of single-unit CP occurrence during maintenance and within-session extinction within the five task windows considered. Given the different number of trials in the two sessions (60 and 69 trials, respectively), we first linearly time-warped extinction trials to fit within 60 bins, which we then used to compute the EDF of that session’s single-unit CPs.

### Onset of behavioral extinction

In order to highlight the neuronal mechanisms initiating behavioral extinction, we chose a drop of 10% in response probability to mark the onset of an extinction-learning episode (behavioral CP_10%_). These drops were defined for each *c*_*b*_ (excluding the single behavioral CP at which response probability increased; **Figure 2A)** as the first trial *t* where 1/(1 + exp(−(*t* − *c*_*b*_)/*τ*_*b*_)) ≥ 0.1 (**Figure 4B**).

### Single-unit and behavioral CP coordination

We collected all single-unit CPs and aligned them with respect to each behavioral CP_10%_ of the corresponding animal (i.e., for animals with two extinction-learning episodes, single-unit CPs were considered twice). The aligned single-unit CPs were then binned using a 3-trial bin. <u>Statistical test:</u> We used permutation bootstraps to test whether single-unit CP frequencies within 10 trials from the behavioral CPs were statistically larger than what is expected by chance. We generated bootstrap histogram samples by randomly shuffling the occurrence of the single-unit CPs of each animal and repeating the alignment procedure described above. This was done by permuting the trial order but keeping co-occurring single-unit CPs of different units at the same trial. Behavioral CPs were left unchanged. The 5000 histogram samples so obtained were then used to produce an EDF over single-unit CP frequency per histogram bin. At each of the 7 frequency values around the behavioral CP of the original single-unit CP histogram a p-value was assigned on the basis of the EDF of their corresponding bin. In order to test significance with a higher temporal resolution than a 3-trial binning, we repeated the bootstrap procedure by sliding the histogram bin edges by 1 and then 2 trials. The p-values assigned to each trial lag (center of the bin) are reported on logarithmic scale in **Figures 4C, 4E** and **S3** as solid black lines. Benjamini-Hochberg’s correction for multiple comparisons was performed only on the p-values of the 7 bins of the displayed histogram. Asterisks mark p<0.05 (black; significant) and p<0.1 (gray; not significant) after correction. <u>Control with maintenance single-unit CPs:</u> To make sure that the results reported in **Figures 4C-E** and **S3B right** are not only due to random PL fluctuations, observed also during maintenance, we repeated the test using behavioral CPs from the extinction session but single-unit CPs from maintenance (**Figure S3A** and **S3B left**).

### Predicting behavioral CPs from PL population vectors

We tested if the firing rate of the PL population in the four task windows examined could predict changes in animal behavior. For each animal, we considered the population vector 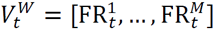 constructed from firing rates 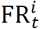 of units *i* in trial *t* in the task window *W*. On the basis of these population vectors, we then trained a support vector machine classifier with linear kernel (slack variables minimized with L^1^ norm and box constraint = 1) to divide the trials occurring before the first behavioral CP from those occurring after the behavioral CP. For this analysis, we only considered the first behavioral CP, in order to have a comparable chance level across animals. Classifier accuracy was computed with a 10-fold cross-validation to avoid overfitting. Since the sample was imbalanced and the two classes (before/after behavioral CP) were not of equal size, we used the Cohen’s kappa coefficient to quantify classifier accuracy relative to chance level. Cohen’s kappa ranges between −1 and 1 and is defined as,

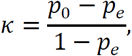

with *p*_0_ the fraction of correctly classified samples and *p*_*e*_ the expected probability of correct classification due to chance. Kappa values were computed on the classifier output collected over the 10 folds for each animal and each task window (**Figures 4F** and **S4C**). <u>Statistical test:</u> Significance of the kappa coefficients was tested through bootstrap. Any monotonic change in firing rate can improve the performance of a classifier trained to divide temporally ordered samples. To account for this factor and test exclusively for the behavioral CP and population-rate coordination, we created a bootstrapped sample by repeatedly assigning the behavioral CP to a random trial. We then trained the classifier to divide trials occurring before the shuffled behavioral CP from those occurring after it. The procedure was repeated 100 times for each animal and each task window. The obtained *κ*^*boot*^ values were then averaged per animal and task window, generating a reference set 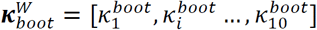 with *i* = 1 … 10 indexing the animal. Since the performance of a classifier highly depends on the number of units composing the population vector, we compared the set of original *κ* values with 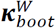 with a one-tailed paired t-test. Significance was assessed with Benjamini-Hochberg’s correction for multiple comparisons.

### Coordination between ITI and CL, DP, and LP single-unit CPs

To quantify coordination between single-unit CPs detected within the ITI window and those of the same unit detected within other windows, we proceeded as follows: For each single-unit CP detected in the ITI window, we computed the distance in trials between its occurrence and that of the nearest single-unit CP of the same unit detected in a different task window, e.g. DP. Absolute distance values were then averaged across all units of an animal (**Figure 4I**).

## ACKNOWLEDGEMENTS

We would like to thank Drs. Ainhoa Bilbao, Thomas Enkel, Wolfgang Kelsch, Andreas Meyer-Lindenberg, Max Scheller, Lennart Oettl and Wolfgang Sommer for their support and stimulating discussions.

## AUTHOR CONTRIBUTION

Investigation, T.M., R.S. and G.K.; Electrophysiology, G.K. and T.M.; Spike Sorting, T.M.; Analysis, Visualization and Interpretation, E.R. and H.T.; Behavioral Model Development, H.T.; Writing - Original Draft, E.R., H.T. and G.K.; Writing - Review and Editing, R.S. and D.D.; Funding Acquisition, T.M., G.K., D.D. and R.S.

## Supplementary Figures

**Figure S1.**
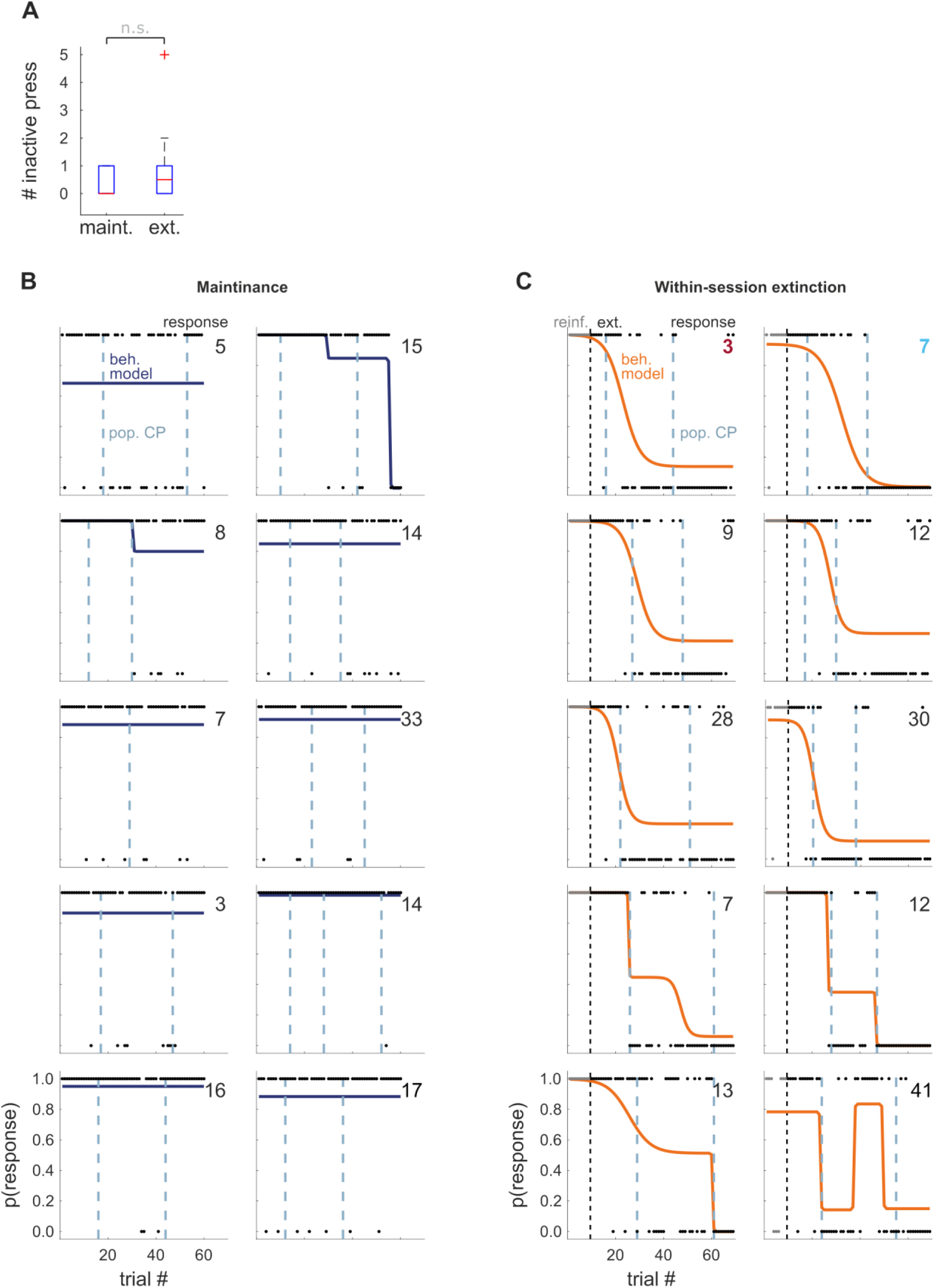
Behavioral and whole-trial population neural changes during maintenance and extinction. (A) Distribution of inactive lever presses per animal during maintenance (left) and extinction (right). See also **Figure 1A**. (B, C) Behavioral models for all animals and their respective whole-trial population CPs during maintenance (B) and extinction (C). Panels at the same relative position in (B) and (C) belong to the same animal. Numbers at the top-right of each panel indicate the number of recorded units. Magenta and cyan in (C) identifies the same animals color-marked in **Figures 2B** and **2C**.

**Figure S2.**
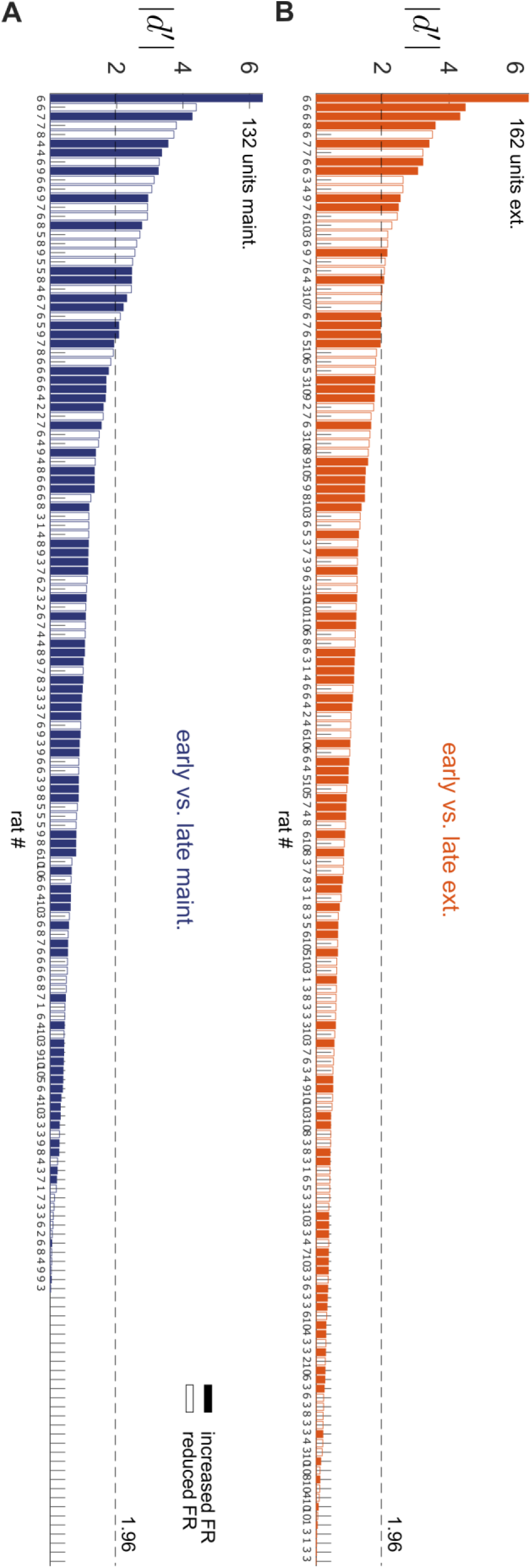
Within-session single-unit rate changes. (A, B) Sensitivity analysis as in **Figure 3H** performed on all recorded units during maintenance (A) and extinction (B). Vertical axis indicates the number of the animal from which the unit was recorded. Dashed lines mark the threshold of significant change in firing rate in units of standard deviations.

**Figure S3.**
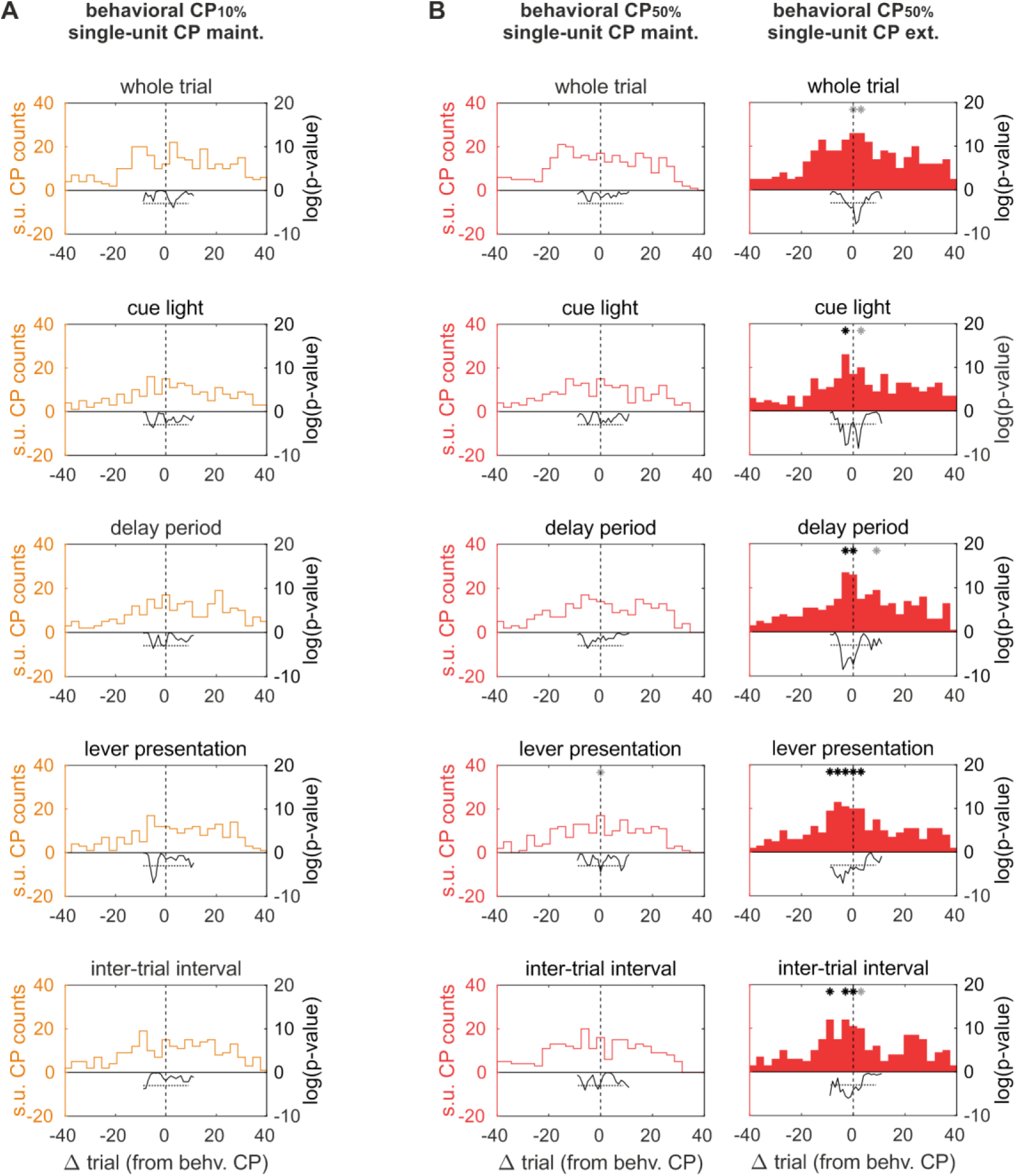
Single-unit CPs anticipate behavioral extinction. (A) Aligning maintenance single-unit CPs to extinction behavioral CPs_10%_. Coordination is lost when single-unit CPs and behavioral CPs belong to different sessions. See also **Figures 4C-E**. (B) Aligning maintenance (left) or extinction (right) single-unit CPs to extinction behavioral CPs_50%_. Coordination is lost when single-unit CPs and behavioral CPs belong to different sessions. See also **Figures 4C-E**.

**Figure S4.**
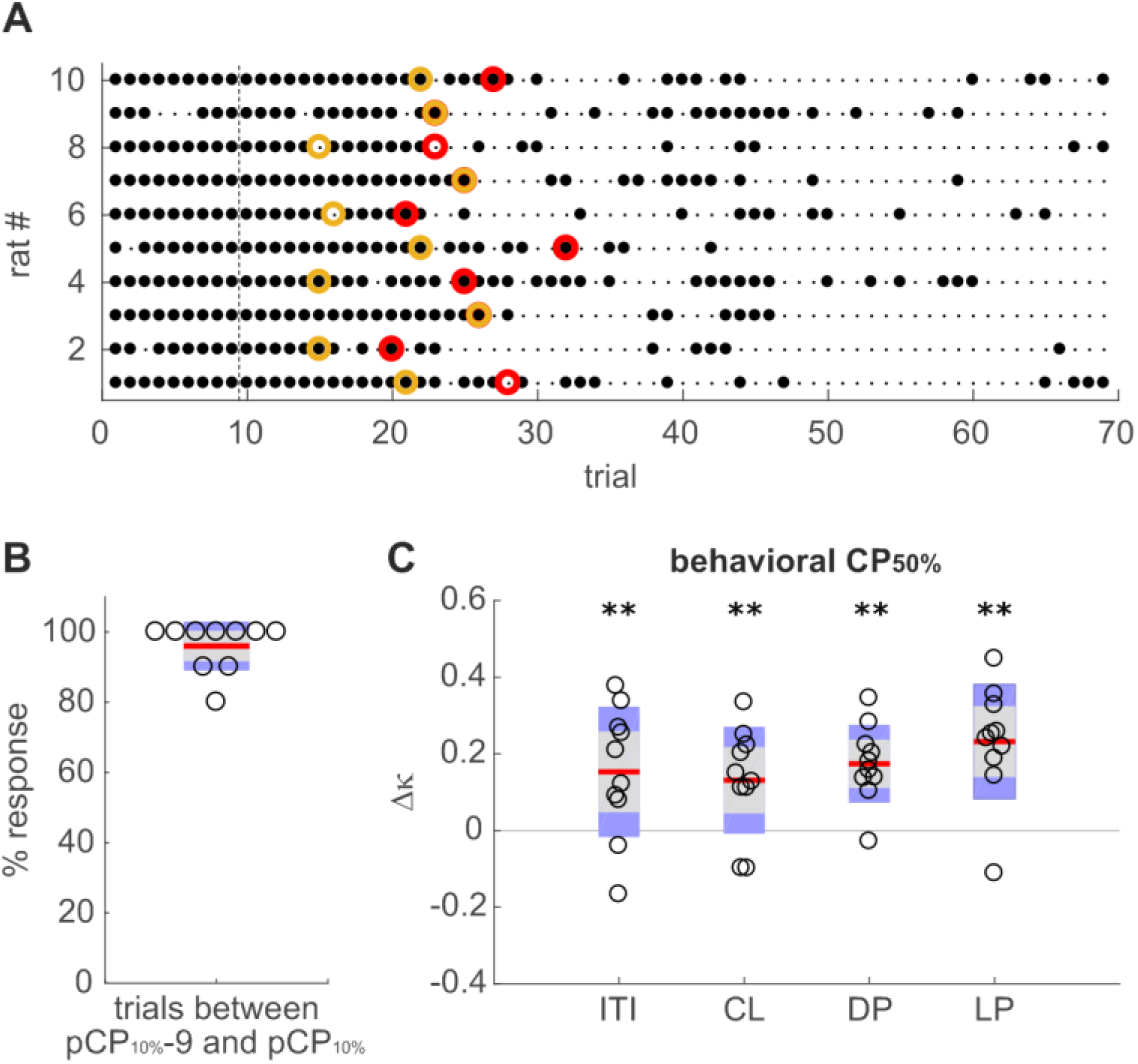
PL population rate is predictive of behavioral extinction. (A) Active lever presses (filled black circles) against first behavioral CP_10%_ (orange) and behavioral CP_50%_ (red) for each animal during extinction. Dashed line indicates the onset of extinction trials. (B) Behavioral CPs_10%_ anticipate behavioral extinction. Open circles show the percentage of active lever presses during the ten trials prior and including behavioral CP_10%_. Plots show mean ± 1.96 · sem (red and gray) and sd (purple). At behavioral CP_10%_ the rats still have not changed their conditioned behavior. (C) Classifier performance as in **Figure 4F** but predicting behavioral CP_50%_. Extinction was predicted form population rates equally well in all task windows.

## REFERENCES

Babayan, B.M., Uchida, N., and Gershman, S.J. (2018). Belief state representation in the dopamine system. Nat Commun 9, 1891.

Benjamini, Y., and Hochberg, Y. (1995). Controlling the false discovery rate: a practical and powerful approach to multiple testing. Journal of the Royal statistical society: series B 57, 289–300.

Botvinick, M., Ritter, S., Wang, J.X., Kurth-Nelson, Z., Blundell, C., and Hassabis, D. (2019). Reinforcement Learning, Fast and Slow. Trends Cogn Sci 23, 408–422.

Brebner, L.S., Ziminski, J.J., Margetts-Smith, G., Sieburg, M.C., Reeve, H.M., Nowotny, T., Hirrlinger, J., Heintz, T.G., Lagnado, L., Kato, S., et al. (2020). The emergence of a stable neuronal ensemble from a wider pool of activated neurons in the dorsal medial prefrontal cortex during appetitive learning in mice. J Neurosci 40, 395–410.

Caballero, J.P., Scarpa, G.B., Remage-Healey, L., and Moorman, D.E. (2019). Differential effects of dorsal and ventral medial prefrontal cortex inactivation during natural reward seeking, extinction, and cue-induced reinstatement. eNeuro 6.

Chen, B.T., Yau, H.J., Hatch, C., Kusumoto-Yoshida, I., Cho, S.L., Hopf, F.W., and Bonci, A. (2013). Rescuing cocaine-induced prefrontal cortex hypoactivity prevents compulsive cocaine seeking. Nature 496, 359–362.

Dunsmoor, J.E., Niv, Y., Daw, N., and Phelps, E.A. (2015). Rethinking extinction. Neuron 88, 47–63.

Durstewitz, D., Vittoz, N.M., Floresco, S.B., and Seamans, J.K. (2010). Abrupt transitions between prefrontal neural ensemble states accompany behavioral transitions during rule learning. Neuron 66, 438–448.

Enel, P., Procyk, E., Quilodran, R., and Dominey, P.F. (2016). Reservoir computing properties of neural dynamics in prefrontal cortex. PLoS Comput Biol 12, e1004967.

Euston, D.R., Gruber, A.J., and McNaughton, B.L. (2012). The role of medial prefrontal cortex in memory and decision making. Neuron 76, 1057–1070.

Fragale, J.E., Khariv, V., Gregor, D.M., Smith, I.M., Jiao, X., Elkabes, S., Servatius, R.J., Pang, K.C., and Beck, K.D. (2016). Dysfunction in amygdala-prefrontal plasticity and extinction-resistant avoidance: A model for anxiety disorder vulnerability. Exp Neurol 275 Pt 1, 59–68.

Gershman, S.J., and Uchida, N. (2019). Believing in dopamine. Nat Rev Neurosci 20, 703–714.

Goldstein, R.Z., and Volkow, N.D. (2011). Dysfunction of the prefrontal cortex in addiction: neuroimaging findings and clinical implications. Nat Rev Neurosci 12, 652–669.

Harris, K.D., and Thiele, A. (2011). Cortical state and attention. Nat Rev Neurosci 12, 509–523.

Heinz, A., Kiefer, F., Smolka, M.N., Endrass, T., Beste, C., Beck, A., Liu, S., Genauck, A., Romund, L., Banaschewski, T., et al. (2020). Addiction Research Consortium: Losing and regaining control over drug intake (ReCoDe)-From trajectories to mechanisms and interventions. Addict Biol 25, e12866.

Hirokawa, J., Vaughan, A., Masset, P., Ott, T., and Kepecs, A. (2019). Frontal cortex neuron types categorically encode single decision variables. Nature 576, 446–451.

Jonkman, S., Mar, A.C., Dickinson, A., Robbins, T.W., and Everitt, B.J. (2009). The rat prelimbic cortex mediates inhibitory response control but not the consolidation of instrumental learning. Behav Neurosci 123, 875–885.

Karlsson, M.P., Tervo, D.G., and Karpova, A.Y. (2012). Network resets in medial prefrontal cortex mark the onset of behavioral uncertainty. Science 338, 135–139.

Katori, Y., Sakamoto, K., Saito, N., Tanji, J., Mushiake, H., and Aihara, K. (2011). Representational switching by dynamical reorganization of attractor structure in a network model of the prefrontal cortex. PLoS Comput Biol 7, e1002266.

Kihlberg, J., Herson, J., and Schotz, W. (1972). Square root transformation revisited. Appl Statist 20, 76–81.

Kim, C.K., Ye, L., Jennings, J.H., Pichamoorthy, N., Tang, D.D., Yoo, A.W., Ramakrishnan, C., and Deisseroth, K. (2017). Molecular and circuit-dynamical identification of top-down neural mechanisms for restraint of reward seeking. Cell 170, 1013–1027 e1014.

Leon, M.I., and Shadlen, M.N. (1999). Effect of expected reward magnitude on the response of neurons in the dorsolateral prefrontal cortex of the macaque. Neuron 24, 415–425.

Ma, L., Hyman, J.M., Durstewitz, D., Phillips, A.G., and Seamans, J.K. (2016). A quantitative analysis of context-dependent remapping of medial frontal cortex neurons and ensembles. J Neurosci 36, 8258–8272.

Mante, V., Sussillo, D., Shenoy, K.V., and Newsome, W.T. (2013). Context-dependent computation by recurrent dynamics in prefrontal cortex. Nature 503, 78–84.

Marek, R., Xu, L., Sullivan, R.K.P., and Sah, P. (2018). Excitatory connections between the prelimbic and infralimbic medial prefrontal cortex show a role for the prelimbic cortex in fear extinction. Nat Neurosci 21, 654–658.

McGinty, V.B., and Grace, A.A. (2008). Selective activation of medial prefrontal-to-accumbens projection neurons by amygdala stimulation and Pavlovian conditioned stimuli. Cereb Cortex 18, 1961–1972.

Mellentin, A.I., Skot, L., Nielsen, B., Schippers, G.M., Nielsen, A.S., Stenager, E., and Juhl, C. (2017). Cue exposure therapy for the treatment of alcohol use disorders: A meta-analytic review. Clin Psychol Rev 57, 195–207.

Moorman, D.E., and Aston-Jones, G. (2015). Prefrontal neurons encode context-based response execution and inhibition in reward seeking and extinction. Proc Natl Acad Sci U S A 112, 9472–9477.

Narayanan, N.S., Kimchi, E.Y., and Laubach, M. (2005). Redundancy and synergy of neuronal ensembles in motor cortex. J Neurosci 25, 4207–4216.

Powell, N.J., and Redish, A.D. (2014). Complex neural codes in rat prelimbic cortex are stable across days on a spatial decision task. Front Behav Neurosci 8, 120.

Powell, N.J., and Redish, A.D. (2016). Representational changes of latent strategies in rat medial prefrontal cortex precede changes in behaviour. Nat Commun 7, 12830.

Puchalla, J.L., Schneidman, E., Harris, R.A., and Berry, M.J. (2005). Redundancy in the population code of the retina. Neuron 46, 493–504.

Quirk, G.J., and Mueller, D. (2008). Neural mechanisms of extinction learning and retrieval. Neuropsychopharmacology 33, 56–72.

Ramanathan, K.R., Jin, J., Giustino, T.F., Payne, M.R., and Maren, S. (2018). Prefrontal projections to the thalamic nucleus reuniens mediate fear extinction. Nat Commun 9, 4527.

Redish, A.D., Jensen, S., Johnson, A., and Kurth-Nelson, Z. (2007). Reconciling reinforcement learning models with behavioral extinction and renewal: implications for addiction, relapse, and problem gambling. Psychol Rev 114, 784–805.

Riaz, S., Puveendrakumaran, P., Khan, D., Yoon, S., Hamel, L., and Ito, R. (2019). Prelimbic and infralimbic cortical inactivations attenuate contextually driven discriminative responding for reward. Sci Rep 9, 3982.

Rich, E.L., and Shapiro, M. (2009). Rat prefrontal cortical neurons selectively code strategy switches. J Neurosci 29, 7208–7219.

Ridderinkhof, K.R., Ullsperger, M., Crone, E.A., and Nieuwenhuis, S. (2004). The role of the medial frontal cortex in cognitive control. Science 306, 443–447.

Schuck, N.W., Gaschler, R., Wenke, D., Heinzle, J., Frensch, P.A., Haynes, J.D., and Reverberi, C. (2015). Medial prefrontal cortex predicts internally driven strategy shifts. Neuron 86, 331–340.

Senn, V., Wolff, S.B., Herry, C., Grenier, F., Ehrlich, I., Grundemann, J., Fadok, J.P., Muller, C., Letzkus, J.J., and Luthi, A. (2014). Long-range connectivity defines behavioral specificity of amygdala neurons. Neuron 81, 428–437.

Sharpe, M.J., Stalnaker, T., Schuck, N.W., Killcross, S., Schoenbaum, G., and Niv, Y. (2019). An integrated model of action selection: distinct modes of cortical control of striatal decision making. Annu Rev Psychol 70, 53–76.

Singh, A., Peyrache, A., and Humphries, M.D. (2019). Medial prefrontal cortex population activity is plastic irrespective of learning. J Neurosci 39, 3470–3483.

Sotres-Bayon, F., Sierra-Mercado, D., Pardilla-Delgado, E., and Quirk, G.J. (2012). Gating of fear in prelimbic cortex by hippocampal and amygdala inputs. Neuron 76, 804–812.

Sparta, D.R., Hovelso, N., Mason, A.O., Kantak, P.A., Ung, R.L., Decot, H.K., and Stuber, G.D. (2014). Activation of prefrontal cortical parvalbumin interneurons facilitates extinction of reward-seeking behavior. J Neurosci 34, 3699–3705.

Stoll, F.M., Fontanier, V., and Procyk, E. (2016). Specific frontal neural dynamics contribute to decisions to check. Nat Commun 7, 11990.

Toutounji, H., and Durstewitz, D. (2018). Detecting multiple change points using adaptive regression splines with application to neural recordings. Front Neuroinform 12, 67.

Toutounji, H., and Pipa, G. (2014). Spatiotemporal computations of an excitable and plastic brain: neuronal plasticity leads to noise-robust and noise-constructive computations. PLoS Comput Biol 10, e1003512.

Voigts, J., Siegle, J.H., Pritchett, D.L., and Moore, C.I. (2013). The flexDrive: an ultra-light implant for optical control and highly parallel chronic recording of neuronal ensembles in freely moving mice. Front Syst Neurosci 7, 8.

Watanabe, M. (1996). Reward expectancy in primate prefrontal neurons. Nature 382, 629–632.

Wimmer, K., Nykamp, D.Q., Constantinidis, C., and Compte, A. (2014). Bump attractor dynamics in prefrontal cortex explains behavioral precision in spatial working memory. Nat Neurosci 17, 431–439.

